# Differentiation for drought strategy but conserved plasticity to heat and drought in locally adapted populations of *Arabidopsis thaliana*

**DOI:** 10.64898/2026.01.07.698234

**Authors:** Sophia F. Buysse, Victoria Nicholes, Jeffrey K. Conner, Emily B. Josephs

## Abstract

**Background and Aims:** Adaptive plasticity may be crucial for plants to survive rapid environmental changes long enough for evolutionary adaptation to occur. Heat and drought are both abiotic stresses projected to increase globally and their individual impact on plant growth and function is well-characterized. However, to understand potential responses to climate change, we must manipulate multiple stressors simultaneously.

**Methods:** We measured heat- and drought-related traits as well as fitness in two locally adapted populations of *Arabidopsis thaliana* from Rödåsen, Sweden and Castelnuovo di Porto, Italy. We used chamber common gardens that simulate the current fall and spring climate in Sweden and a hotter and drier climate to identify population differentiation for trait means, trait plasticity, and fitness.

**Key Results:** Population differentiation in both treatments suggests that the Swedish population avoids drought by investing in stress-tolerant leaves while the Italian population escapes drought by flowering early. Despite these differences, there is little evidence for genetic differentiation of plasticity; when experiencing heat and drought, both populations shift their traits in the direction expected to avoid drought. Further, both populations have greatly decreased fitness in heat and drought.

**Conclusions:** These results highlight that locally adapted populations with genetic differentiation for traits within a single environment can respond to the same stressors with plasticity in the same direction. Further, the combination of heat and drought will be extremely damaging to plant populations.

**Lay Summary:** Plant populations respond to environmental stress in different ways, and their response can determine their survival. We found that plant populations that have evolved different drought responses in their native environments still have the same drought response in an experimental hot and dry environment. Both populations also make many fewer fruits, indicating they may decline in future climates.

## INTRODUCTION

Plants need to respond through plasticity or genetic change in order to persist during rapid environmental shifts (Franks *et al*. 2014). Plasticity is the ability of a genotype to produce different phenotypes in response to different environmental conditions (Bradshaw 1965). Adaptive plasticity allows individuals to change their phenotype to move toward the fitness optimum in different environments (Dudley and Schmitt 1996; Price *et al*. 2003; Ghalambor *et al*. 2007; Bell and Galloway 2008). Plasticity is expected to evolve in temporally variable environments and in spatially variable environments connected by gene flow (Bradshaw 1965; Scheiner 1998; Sultan and Spencer 2002; Van Buskirk 2002; Kawecki and Ebert 2004). Adaptively plastic populations are more likely to survive rapid environmental changes through an immediate plastic response and persist long enough to genetically adapt to new environments (Bradshaw 1965; Price *et al*. 2003, 2008; Yeh and Price 2004; Ghalambor *et al*. 2007; Handelsman *et al*. 2013; Morris 2014; Diamond and Martin 2021; Pfennig 2021).

How plants respond to rapid environmental change will depend on how their past evolutionary history has shaped both trait means and trait plasticity. Within-species and within-population genetic variation for trait means in a single environment and for trait plasticity between environments is common and has been frequently documented in plants (Dudley and Schmitt 1996; Gianoli and González-Teuber 2005; Bell and Galloway 2008; Murren *et al*. 2014). As a result, exposing plant populations to environments that simulate combinations of abiotic stresses projected to occur in future climates is crucial to predict adaptation and inform conservation (Hamann *et al*. 2021; Zandalinas and Mittler 2022). We are particularly interested in plasticity to heat and drought because both are projected to increase globally in addition to changes in climate variability (IPCC 2021).

Escape, avoidance, and tolerance are well-studied strategies for adapting to drought stress (Ludlow 1989) but their specific definitions can vary between studies. Plants using the escape strategy have rapid growth to set seed before a drought is lethal; they are found in locations where a drought ends the growing season. Traits associated with escape include early flowering, low water use efficiency (i.e., use more water in biomass production), and minimal investment in protective metabolites and proteins (Ludlow 1989; Kenney *et al*. 2014). If caught in a drought, escape plants show signs of stress including wilting or decreased relative water content (Blum 2014). In contrast, avoidant plants experience drought during their life cycle, rather than at the end of the growing season, and maintain their leaf relative water content to avoid wilting (Ludlow 1989). Traits associated with avoidance include slow growth, later flowering, high water use efficiency, and increased root-to-shoot biomass ratio. Avoidant plants invest in stress protective pathways to allow metabolic and stomatal adjustments which can be reflected in leaves with low specific leaf area and high leaf dry matter content (Ludlow 1989; Des Marais *et al*. 2012; Kenney *et al*. 2014; Kooyers 2015). Tolerant plants have tissues that can survive periods of severe dehydration and remain functional during severe water reduction, such as resurrection plants (Ludlow 1989). We follow these descriptions and define escape plants as quickly flowering to finish their life cycle before the onset of drought, avoidant plants as slow growing to maintain high relative water content, and tolerant plants as continuing to function at extremely low water content.

Plants may have traits associated with more than one drought strategy (Levitt 1980; Ludlow 1989). While drought strategies are typically used to describe differences between species, variation in drought strategy can occur between populations of the same species. For example, *Arabidopsis thaliana* is generally drought avoidant but some populations from across the European range flower early and have lower water use efficiency consistent with drought escape (Mckay *et al*. 2003; Verslues and Juenger 2011; Lovell *et al*. 2013; Kenney *et al*. 2014). Prior work identified *FRI* as a pleiotropic gene involved in variation between drought escape and avoidance in *A. thaliana* where low expression *FRI* alleles are associated with early flowering, low water use efficiency, and faster growth while high expression is associated with later flowering, higher water use efficiency and slow growth (Mckay *et al*. 2003; Juenger *et al*. 2005; Lovell *et al*. 2013).

Understanding a population’s plastic response to future climates requires experimentally manipulating multiple abiotic stressors in tandem, particularly heat and drought (Anderson 2016). Heat is a well-characterized abiotic stress that in *A. thaliana* decreases growth, increases specific leaf area (SLA), and decreases stomatal density among many physiological and molecular responses (Vasseur *et al*. 2011; Vile *et al*. 2012; Suter and Widmer 2013; Stewart *et al*. 2016; Gao *et al*. 2020). Heat can exacerbate the negative effect of drought (Zhang and Sonnewald 2017; Schepers *et al*. 2024); in *A. thaliana,* the rate of leaf production decreased 16% in heat and 23% in drought but 40% when heat and drought were experienced together (Vile *et al*. 2012). However, for traits such as SLA, root-to-shoot ratio, and stomatal density, plastic responses to drought and heat are in opposite directions that cancel out when heat and drought are experienced together (Vile *et al*. 2012; Zhang and Sonnewald 2017). The interaction of heat and drought, whether their individual effects are in the same or opposite directions, may determine population survival in future climates. Many populations of *A. thaliana* are winter annuals that emerge in the fall, overwinter as rosettes, and flower in the spring. Increased spring temperatures may exacerbate drought conditions during flowering for winter annual plants (IPCC 2021). Although earlier arrival of spring may select for earlier flowering and favor low-expression *FRI* alleles, this may come at the cost of reduced water use efficiency during increasingly frequent drought prior to flowering because of the pleiotropic effects of *FRI*. This potential temperature shift in the growing season could mean that more southern populations already adapted to higher temperatures may be better adapted to future conditions at the northern edge of the range than the populations currently growing there (Wilczek *et al*. 2014).

We use two locally adapted populations of *A. thaliana* from Rödåsen, Sweden and Castelnuovo di Porto, Italy (hereafter "Sweden" and "Italy"; Ågren & Schemske, 2012) to look for genetic differentiation in traits indicative of differences in drought strategy. We then test for differentiation for plasticity in response to heat and drought and identify how heat and drought impact fitness in each population. Both populations are winter annuals that emerge in the fall, overwinter as rosettes, and flower in the spring (Ågren and Schemske 2012). The climate at both field sites is remarkably different with a long, cold winter at the Swedish site and a Mediterranean climate at the Italian site with a cool, wet winter but a very hot and dry summer. More specifically, the Swedish field site has less precipitation and more variation in water availability than the Italian field site between germination and flowering (Mojica *et al*. 2016).

The temperature at both sites is similar during germination and flowering, but the Swedish site is much colder during the intermediate months; the soil temperature at the Swedish field site is below freezing for multiple weeks in the winter (Dittmar *et al*. 2014; Mojica *et al*. 2016; Durán *et al*. 2022). In contrast, the Italian field site is hotter and drier than the Swedish site in the summer months, typically at the end of flowering, and the Italian population produces seeds with strong dormancy to survive in the seed bank until conditions are favorable (Postma *et al*. 2016). In both reciprocal transplant studies and chamber common gardens, the Swedish population avoids drought, indicated by higher water use efficiency, and flowers later than the Italian population (Ågren and Schemske 2012; Ågren *et al*. 2013; Dittmar *et al*. 2014; Mojica *et al*. 2016). The Swedish population is also generally more plastic in response to experimental drought and heat (Stewart *et al*. 2016; Mojica *et al*. 2016). Conversely, the Italian population is adapted to a mild winter with fewer calendar days between germination and flowering; it flowers early to escape the severe drought during the hot Italian summer (Ågren and Schemske 2012; Ågren *et al*. 2013; Dittmar *et al*. 2014; Royer *et al*. 2016; Stewart *et al*. 2016). However, much of the prior work on these populations uses only one genotype per population (Cohu *et al*. 2013; Stewart *et al*. 2015, 2016; Mojica *et al*. 2016; Durán *et al*. 2022) and/or investigates seed dormancy, cold tolerance, germination timing, flowering time or fitness (Ågren and Schemske 2012; Ågren *et al*. 2013, 2017; Grillo *et al*. 2013; Oakley *et al*. 2014, 2015, 2018, 2023; Dittmar *et al*. 2014; Postma *et al*. 2016; Postma and Ågren 2016, 2018; Price *et al*. 2018; Sanderson *et al*. 2020; Zacchello *et al*. 2020; Ellis *et al*. 2021; Lee *et al*. 2024; Ellis and Ågren 2024). We build on this work by evaluating plasticity in a range of heat- and drought-related traits (days to flowering, relative water content, specific leaf area, stomatal density, and biomass allocation) to a simulated hot and dry future climate and measuring fitness of multiple genotypes per population.

We grew native genotypes from Sweden and Italy in two chamber common garden experiments that each simulated the current fall and spring climate in Sweden and a hotter and drier potential future Swedish climate. We investigated the following questions:

1. When drought is combined with heat, does genetic differentiation of heat- and drought-related traits support that Sweden avoids drought and Italy escapes drought as described by previous drought studies?
2. Assuming Sweden avoids drought and Italy escapes drought, is there genetic differentiation for plasticity to heat and drought between these two populations that evolved in climates that differ in heat and drought variability?
3. How do heat and drought influence differences in fitness between locally adapted populations in our simulated potential future Swedish climate?

We found consistent genetic differentiation for drought strategy where Italy escapes and Sweden avoids when heat and drought are experienced together. Despite this, both populations had plasticity toward drought avoidance at a similar magnitude for almost all traits. There was no difference in fitness between the populations within each treatment, likely due to our lack of freezing temperatures during the simulated winter, but both populations greatly decreased fitness in our simulated future climate. Despite difficulties simulating Swedish winter in our chamber common gardens, this raises concerns about population persistence in future climates.

## MATERIALS AND METHODS

### 2021 Experiment

#### Seed Material and Growing Conditions

Seeds from Rödåsen, Sweden and Castelnuovo di Porto, Italy were collected in the field (Ågren and Schemske 2012). Native genotypes, including the parents of a set of recombinant inbred lines (Ågren *et al*. 2013), were grown in chamber common gardens for at least one generation. We sowed seeds (3-5 per pot) from eight Italian genotypes and 13 Swedish genotypes directly on wet soil and stratified them at 4°C for five days in the dark. After stratification, pots were split into two common gardens in BioChambers growth chambers at Michigan State University, East Lansing, MI. One common garden simulated “Current” climate conditions near the Sweden field collection site in the fall and spring and the other simulated a “Future” hotter and drier climate at the same location. Each treatment included six replicates of the recombinant inbred line parents and two replicates of each of the remaining seven genotypes from Italy and 12 genotypes from Sweden [(6 replicates x 2 RIL parents) + (2 replicates x 7 Italy lines) + (2 replicates x 12 Sweden lines) x 2 treatments = 100 plants]. Plants were grown in a soil mix of two parts Michigan Grower Products SureMix (Michigan Grower Products, 251 McCollum, Galesburg, MI 49053) to one part Sunshine Redi-Earth (Sun Gro Horticulture, 770 Silver Stret, Agawam, MA 01001) supplemented with Osmocote 15-9-12 slow-release fertilizer (1 tablespoon fertilizer was split evenly between 25 pots; The Scotts Company, 14111 Scottslawn Road, Marysville, OH 43041).

Temperatures in each treatment were determined using data from Weather Underground (weatherunderground.com) spanning 1998 to 2020. Median day and night air temperatures from the Sundsvall-Timra Airport (49km from the Sweden field collection site) were used to determine the “Current” treatment conditions. In the field, plants emerge in August and September (Ågren and Schemske 2012). Accordingly, we split temperatures from August 1^st^ through October 31^st^ into four chronological subsets and used the median day and night temperature for each subset as the chamber temperatures for one week each (Supplementary Table 1). Daylength was determined by following the same procedure. By the end of October, the mean temperature at the Sweden field site is below 10°C which is the minimum for our chambers. At this point, plants were thinned to one per pot and vernalized for five weeks in a cold room where temperatures varied from 7-10°C during the day and were 8°C at night. Our cold treatment is sufficient to induce flowering in both populations though it is much warmer than temperatures at the Sweden field site during winter (Dittmar *et al*. 2014; Mojica *et al*. 2016; Zacchello *et al*. 2020; Lee *et al*. 2024). The daylength for each week of our simulated winter was set by splitting daylengths from November through April into five subsets and using the median from each subset.

Plants were then returned to growth chambers to simulate spring. At the Sweden field site, plants flower in May and June (Ågren and Schemske 2012). We simulated spring using the same approach of the median day temperature, night temperature, and daylength but time was not condensed. In the spring simulation, one week in the experiment corresponded to one week of temperatures from the field to allow plants time to flower. Flowering began while we were simulating June which is near the end of flowering in the field (Ågren and Schemske 2012). Through our experimental approach, we condensed approximately 48 weeks in the field (August through June) into 25 weeks in chambers (Table S1). The “Future” treatment was 4°C hotter than the “Current” treatment when simulating fall and spring following IPCC projections for Sweden within the next 50 years if global temperatures rise by 2°C (IPCC 2021). Winter temperatures were the same between both treatments because our minimum temperature in the Current treatment is already much warmer than the average winter temperatures in the field. While our “Current” treatment does not accurately simulate the current winter climate at the Swedish field site, we use the Current and Future nomenclature to encompass the general trend of a hotter and drier climate that we simulate in the fall and spring.

To manipulate water availability, we added 50g of our soil mix to each pot in the experiment and to 10 additional pots. The additional pots were watered to saturation to determine water holding capacity (100% gravimetric water content) and oven dried at 60°C for one week to determine dried mass (0% gravimetric water content). We used the average 0% and 100% masses from the test pots to calculate the weight required to maintain pots with plants in our treatments at their target gravimetric water content. All pots were well watered when seeds were sowed. The “Current” treatment was then kept at 45% gravimetric water content throughout the experiment to maintain a well-watered environment (Vile *et al*. 2012). Plants in the “Future” treatment were dried down to 25% gravimetric water content before vernalization (Vile *et al*. 2012). Because plants showed no visible signs of water stress by the end of vernalization, they were dried down to 10% gravimetric water content for 3 weeks and further dried to 7.5% gravimetric water content for the final 12 weeks of the experiment. Soil moisture was maintained by daily weighing and watering to account for differences in water use by each plant (Granier *et al*. 2006). Plants were rotated within each chamber once a week to reduce microclimate differences.

#### Phenotyping

We recorded emergence and flowering as phenology timepoints defined as the first day green growth was visible and the first day that white petal was visible from a bud, respectively, after sowing. On the first day of flowering, the last fully developed rosette leaf was collected, typically from the third whorl of leaves, and measured for relative water content, specific leaf area (SLA) and leaf dry matter content as related measures of the leaf economics spectrum (Pérez-Harguindeguy *et al*. 2013). The leaf was weighed immediately after collection (fresh mass: *M*_f_) and scanned to measure leaf area (*A)* with ImageJ (Schneider *et al*. 2012). Each leaf was then rehydrated by wrapping in a wet paper towel and tin foil and placed in the dark at 4°C for 24 to 48 hours. Rehydrated leaves were weighed (*M*_r_) and dried at 45°C for at least two weeks before recording a dry mass (*M*_d_). From these values, we calculated relative water content ((*M*_f_ – *M*_d_)/( *M*_r_ – *M*_d_)), SLA (*A* / *M*_d_), and leaf dry matter content (*M*_d_ / *M*_r_). Relative water content is our measurement of physiological drought stress where a decrease in relative water content in the Future treatment indicates successful drought stress (VanBuren *et al*. 2024).

Plants were destructively harvested in the final two weeks of the experiment when they fit any of the following criteria: 1) three fruits dehisced on the main flowering stalk and few open flowers or buds remain; 2) two fruits dehisced on main flowering stalk and most fruits on branches dehisced; 3) no remaining open flowers or green buds, even if no fruits dehisced; 4) 175 days post planting (25 weeks; simulating the end of July in the field) if they had not met the earlier criteria. Only five of the 81 plants that survived were harvested following the final criterion; all were Sweden genotypes and four were in the Current treatment. At harvest, we counted rosette leaf number, and dried rosette and reproductive biomass at 45°C for at least two weeks to analyze reproductive allocation. We manually counted fruit number in ImageJ with photos of all plant branches and collected ten fruits from the main flowering stalk of each plant to calculate the average seed weight and the average number of seeds per fruit. To measure total fitness, the total number of fruits produced by a plant was multiplied by the average number of seeds per fruit. This is an unusually complete measure of total male and female fitness because *A. thaliana* is 95% self-fertilizing (Abbott and Gomes 1989; Platt *et al*. 2010).

Plasticity in response to the Future treatment was measured by subtracting the mean trait value for a given genotype in the Current treatment from that genotype’s mean trait value in the Future treatment.

#### Statistical Analysis

All statistical analyses were done in R (v. 4.4.1; R Core Team 2024) with the following packages: *tidyverse*, *lme4*, *lmerTest*, and *emmeans* (Bates *et al*. 2015; Kuznetsova *et al*. 2017; Wickham *et al*. 2019; Lenth 2024). The Current and Future treatments were analyzed using type III ANOVAs (Des Marais *et al*. 2013; Laitinen and Nikoloski 2019) with treatment, population, and their interaction as fixed effects and genotype nested within population as a random effect. The treatment fixed effect tests for plasticity across both populations, the population fixed effect tests for population differentiation for trait means across the treatments, and their interaction tests for population differentiation for plasticity. SLA and average seed number per fruit were log transformed to increase the normality of model residuals. Model predicted means and 95% confidence intervals were plotted with *ggplot* (Wickham 2016); log-transformed traits were back transformed for plotting.

### 2022 Experiment

#### Seed Material and Growing Conditions

We conducted a follow-up experiment in April 2022 using similar methods but focusing on allocation to above and below ground biomass at bolting instead of lifetime fitness. Root-to-shoot biomass ratio increases under severe drought to access additional soil water and reduce leaf transpiration (Verslues and Juenger 2011). Eight pots were directly sown with seeds from each of the eight Italian genotypes and eight randomly chosen Swedish genotypes that were grown in the 2021 experiment. Four replicates of each genotype were planted in each treatment (16 genotypes x 4 replicates x 2 treatments = 128 plants). Plants were grown in Sunshine Redi-Earth amended with 3.6kg/m^3^ 15-9-12 Osmocote slow-release fertilizer. Soil mass, water holding capacity and dried mass were determined as in the 2021 experiment. Seeds were stratified in the dark at 8°C for 5 days before being moved to a BioChambers growth chamber at Michigan State University, East Lansing, MI. Due to time constraints, all pots were grown in 16°C day temperatures, 13°C night temperatures with 16-hour days for three weeks (Table S2). Plants were then thinned to one per pot or transplanted to new pots to replace low survival of certain genotypes (Italy only). After thinning and transplanting, each population had the same number of plants, but some Italian genotypes had more than four replicates per treatment and some had fewer (min = 1, max = 6; Current treatment mean = 4.57, Future treatment mean = 4). All pots were vernalized as in the 2021 experiment (Table S1). After vernalization, plants were moved to Percival Growth Chambers with photosynthetically active radiation of 88-90 µmol/m2/s at the Kellogg Biological Station, Hickory Corners, MI. Plants were randomized within 2 blocks for each treatment, and each block was grown in a different growth chamber. Chambers simulated spring using the same temperatures and daylengths from the 2021 experiment (Table S2). In the 2021 experiment, we noticed that light levels were lower in the Current treatment which had lower temperatures. To remedy this, in 2022 shelf heights within the chamber were adjusted each week as temperatures were changed to ensure a consistent 88-90 µmol/m2/s of light at the soil surface and plants were rotated weekly to reduce microclimate differences. We focus our conclusions on results shared between both experiments that are likely from heat and drought rather than results that only occurred in 2021 potentially from differences in light.

As in 2021, the Current treatment was consistently maintained at 45% soil moisture. The Future treatment began at 7.5% soil moisture which was increased to 10% in week 14 due to observed wilting and drying of rosette leaves. Plants were weighed and watered daily during the first three weeks of the experiment, twice a week during vernalization, and daily during the eight weeks of spring simulation.

#### Phenotyping

Emergence, rosette leaf number, SLA, leaf dry matter content, and relative water content were measured as in the 2021 experiment with the following exceptions: leaf traits were measured at bolting, the day buds were visible in the center of the rosette, rather than at flowering and biomass was dried at 60°C for at least 48 hours before weighing. A leaf in the third or fourth whorl from the apical meristem was selected for a stomatal peel on the abaxial side of the leaf to measure stomatal density at bolting. Stomata density is likely locally adapted in *A. thaliana* though it is not correlated with water use efficiency (Dittberner *et al*. 2018; de Ollas *et al*. 2023). The whorl selected depended upon the size of the plant as small leaves do not peel cleanly. Two images were taken at 400x magnification for each stomatal peel; each field of view was 0.06612mm^2^. Images were taken at randomly chosen spots on the peels but avoiding overlapping windows. Only stomata entirely in the field of view of the image were counted in ImageJ to determine stomatal density.

Plants were destructively harvested at bolting, or as soon as possible after, to obtain root and shoot biomass (range 0 – 10 days after bolting; median = 4 days). Shoot biomass was collected by cutting plants at the soil surface. Each pot was then placed in a 1% Liquinox solution to soak for 20 minutes. Soil was gently washed away with running water until roots were clean or 30 minutes had passed; most roots were clean within the 30-minute timeframe. Root and shoot biomass was dried at 60°C for 3 days and weighed.

#### Statistical Analysis

All analyses were conducted as in the 2021 experiment with the addition of block nested in treatment as a fixed effect in the model. Stomatal density, SLA, shoot biomass, and root-to-shoot ratio were log transformed to increase normality of residuals before analysis. Model output was back-transformed before plotting for comparison across traits and studies.

## RESULTS

Our heat and drought treatment successfully induced non-lethal drought stress as leaf relative water content decreased at bolting (2022) and flowering (2021). Relative water content decreased in both populations in the Future treatment in both experiments, and more so in 2022; population mean relative water content decreased by 8% (Italian population, 2021), 10% (Italian population, 2022), 5% (Swedish population, 2021), and 7% (Swedish population, 2022) (Fig 1A; 2021 Treatment *p* = 1.50×10^-10^; 2022 Treatment *p* = 7.97×10^-15^).

**Figure 1.**
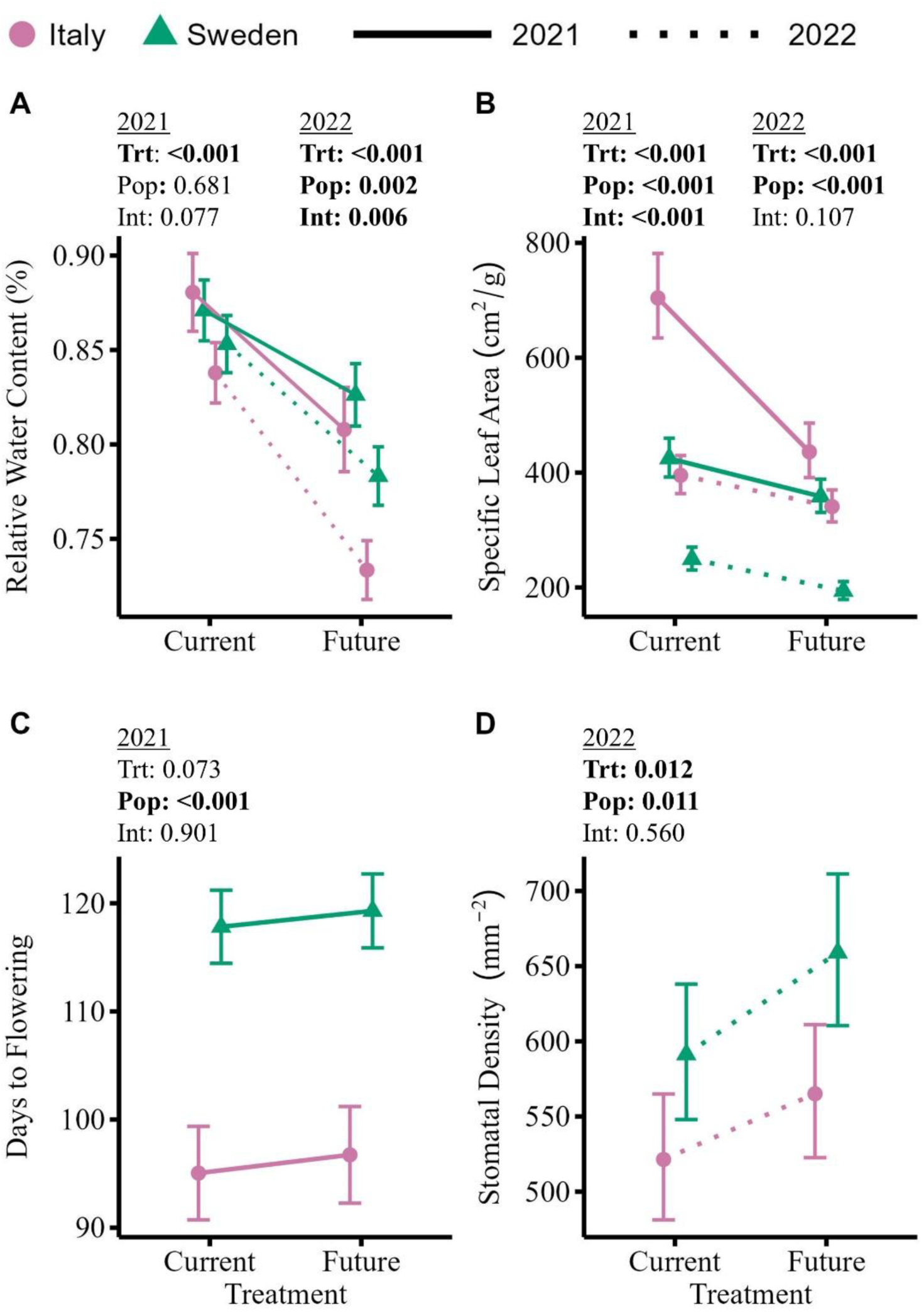
Sweden avoids drought and Italy escapes drought. Each panel displays estimated marginal means for Sweden (triangles) and Italy (circles) with 95% confidence intervals. Lines connecting means illustrate plasticity; the 2021 experiment is connected with solid lines and the 2022 experiment with dotted lines. Statistics above each panel show *p*-values for Anova fixed effects. **Alt Text:** Reaction norms labelled A through D comparing plasticity in relative water content, specific leaf area, days to flowering, and stomatal density between populations and treatments for both experiments. For both populations and experiments, going from the current treatment to the future treatment, relative water content and specific leaf area decrease while days to flowering and stomatal density increase.

### Sweden avoids, Italy escapes

Consistent with genetic differentiation for drought avoidance, the Swedish population had higher relative water content in the Future treatment though the difference is only significant in 2022 (Fig 1A; 2021 Pop *p* = 0.68; 2022 Pop *p* = 0.002). The Swedish population had on average 34% lower SLA (Fig 1B; 2021 Pop *p* = 2.12×10^-5^; 2022 Pop *p* = 2.39×10^-8^), 16% higher leaf dry matter content (Fig S1A; 2021 Pop *p* = 0.028; 2022 Pop *p* = 6.52×10^-10^), and flowered 22 days later (Fig 1C; 2021 Pop *p* = 5.82×10^-7^) than Italy. SLA and leaf dry matter content are highly negatively correlated across all plants because *M_d_* is the denominator of SLA and numerator of leaf dry matter content (2021 *r* = −0.701, *p* <0.001; 2022 *r* = −0.746, *p* < 0.001). Sweden also had 13% (Current) to 16% (Future) greater stomatal density than Italy (Fig 1D; 2022 Pop *p* = 0.011).

### There was little genetic differentiation for plasticity

Despite genetic differentiation for traits within each treatment, there was limited genetic differentiation for plasticity (i.e., the interaction was only significant in one year for relative water content, SLA, root-to-shoot ratio, and shoot biomass; Table 1). Both populations were plastic in the direction expected of drought avoidance; for example, SLA decreased in the Future treatment (Fig 1B; 2021 Trt *p =* 3.76×10^-13^; 2022 Trt *p* = 2.60×10^-9^).

**Table 1.**
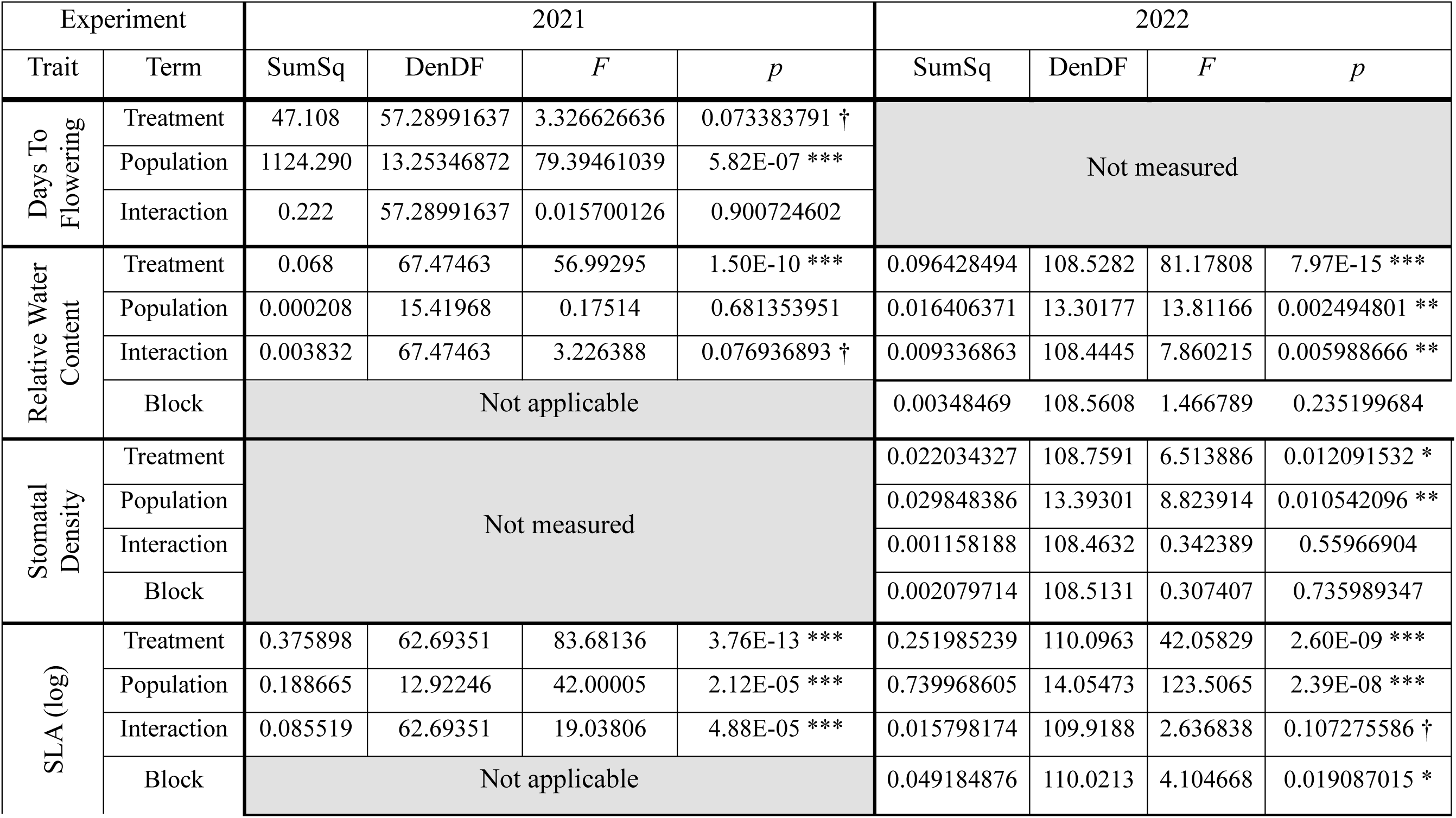
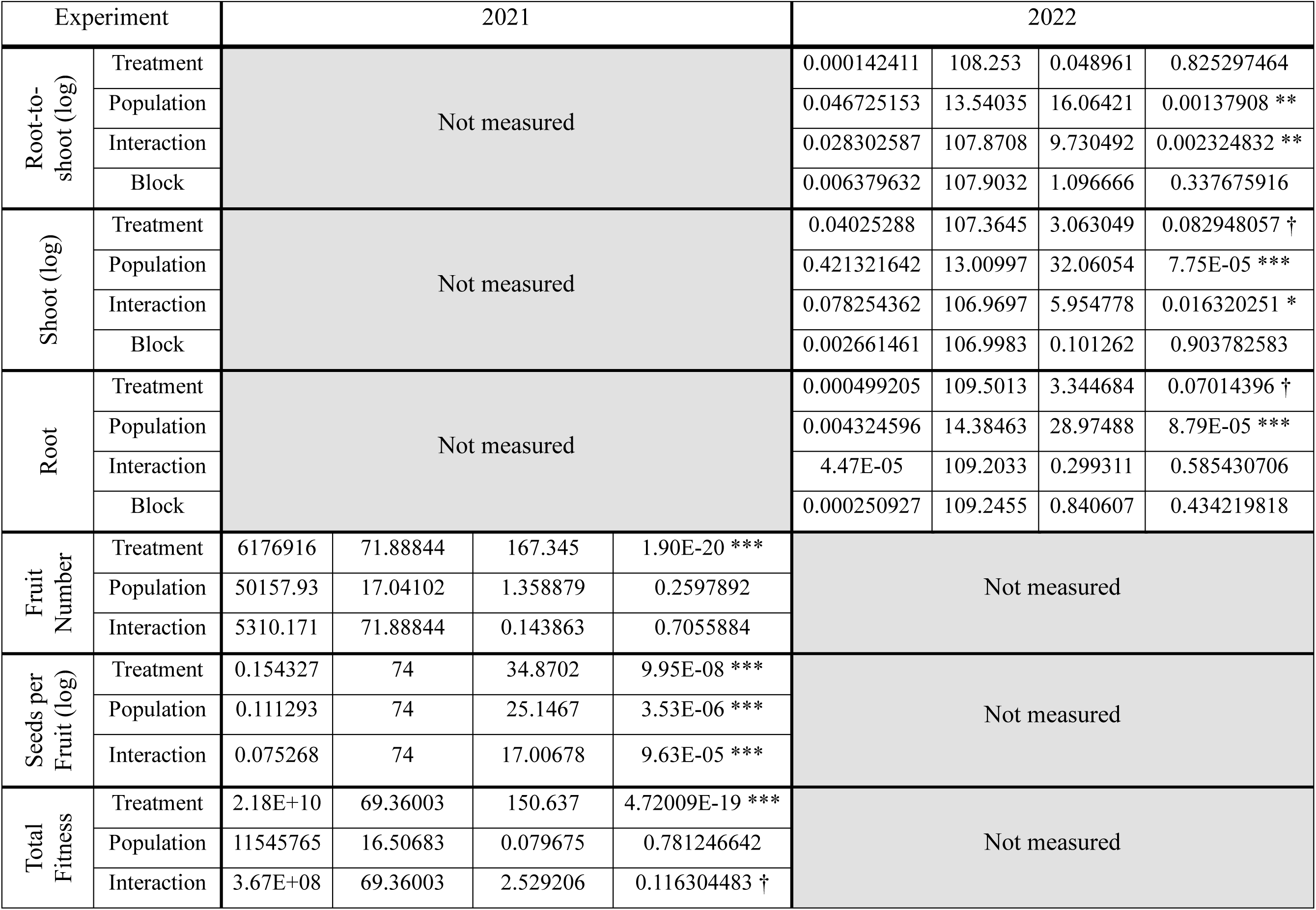
Anova results. Columns denote the trait used as the response variable in the model, the predictor model terms, and then for the 2021 and 2022 experiments the sum of squares (SumSq), the denominator degrees of freedom (DenDF), *F* value, and *p* value. Numerator degrees of freedom is always 1 for treatment, population, and their interaction; it is always 2 for block. Symbols illustrate *p* value: † < 0.1; * <0.05; ** < 0.01; *** < 0.001.

There was evidence for differentiation for plasticity in the components of our leaf traits but neither population was consistently more plastic (Fig S1; Table S3). Thus, the genetic differentiation for plasticity was dependent on the slight difference in experiment conditions between years and genetic differentiation for differing drought strategies did not include genetic differentiation for plasticity in either year.

While escape and avoidance are useful terms to understand plant drought survival, traits associated with each strategy are not prescriptive of all plant traits or trait plasticity. The Italian population had a greater root-to-shoot ratio than the Swedish population (Pop *p* = 0.001) and increased root-to-shoot ratio in the Future treatment, typical of drought avoidance rather than escape, whereas the ratio decreased in the Swedish population (Fig 2A; Verslues & Juenger, 2011). The different directions of plasticity in root-to-shoot ratio between the populations were caused by different patterns of plasticity in the component biomasses. Both populations had lower root biomass in the Future treatment (Fig 2C). The Swedish population maintained consistent shoot biomass across treatments, (Fig 2B) resulting in a decrease in root-to-shoot ratio. In contrast, the Italian population decreased shoot biomass in the Future treatment (Fig 2B). Since the values for shoot biomass were an order of magnitude higher than root biomass, this caused an increase in root-to-shoot ratio. The lower root and shoot biomass of the Italian plants and their plastic decrease in the Future treatment is consistent with drought escape by growing quickly and not investing in stress avoidance despite the observed increase in root-to-shoot biomass.

**Figure 2.**
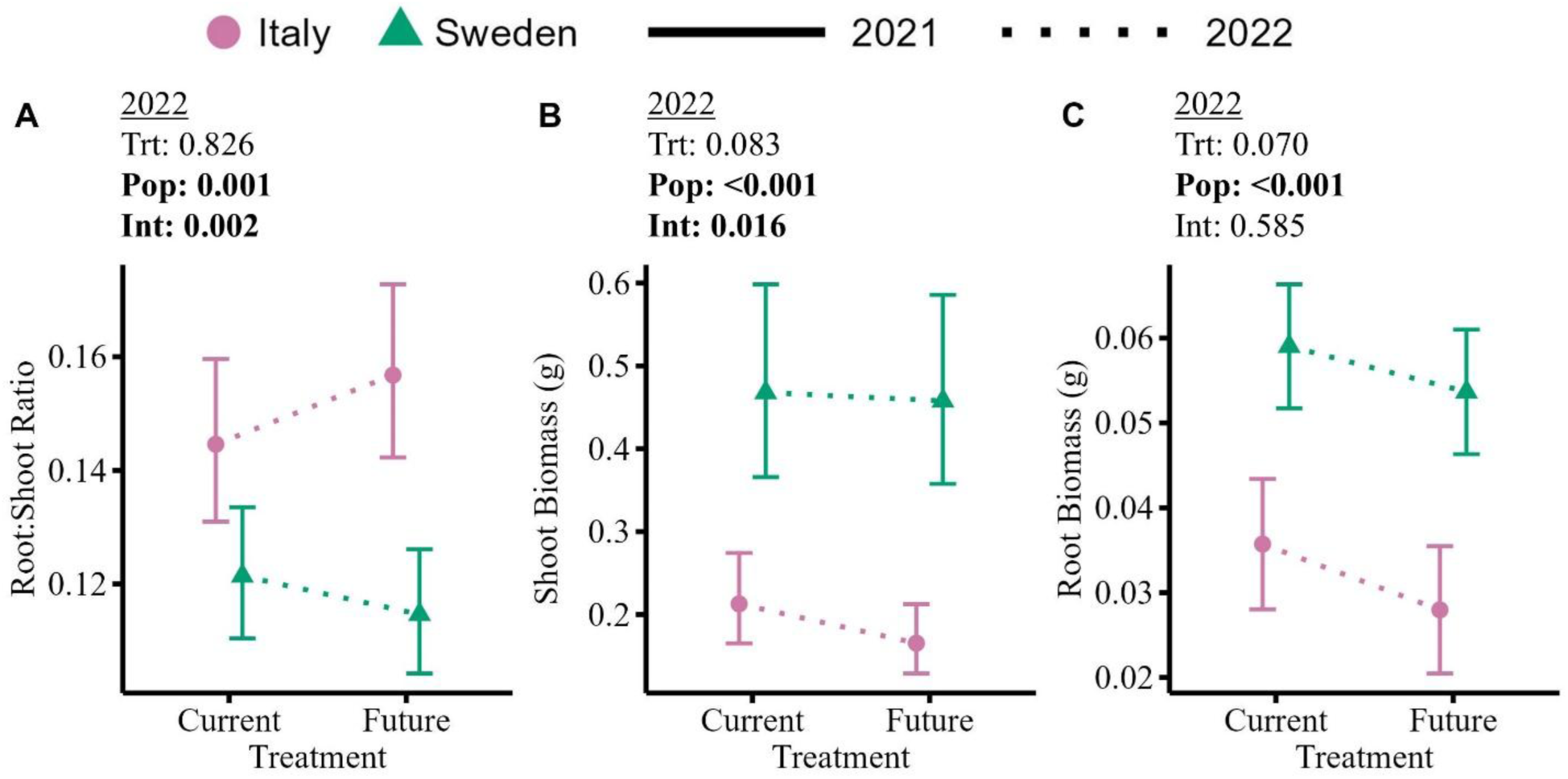
Population genetic differentiation for plasticity in root-to-shoot biomass ratio is driven by a greater decrease in shoot biomass than root biomass in Italy. Each panel displays estimated marginal means for Sweden (triangles) and Italy (circles) with 95% confidence intervals. Lines connecting means illustrate plasticity; these traits were only measured in 2022 (dotted lines). Statistics above each panel show *p*-values for Anova fixed effects. **Alt Text:** Reaction norms labelled A through C comparing plasticity in root to shoot biomass ratio, shoot biomass, and root biomass from 2022. For both populations, going from the current treatment to the future treatment, shoot biomass and root biomass decrease. For Italy, root to shoot biomass ratio increases but it decreases for Sweden.

### Fitness is greatly reduced fitness in the Future treatment

Fruit number (Trt *p* = 1.90×10^-20^) and total fitness (Trt *p* = 4.72×10^-19^) were greatly reduced in both populations in the Future treatment. Fruit number decreased by 78% in the Italian population and 85% in the Swedish population. Total fitness decreased by 80% in the Italian population and 90% in the Swedish population. However, the average number of seeds per fruit decreased significantly only for Sweden (Fig 3; Trt *p* = 9.95×10^-8^; decreased 30% and 6% in the Swedish and Italian populations respectively). Average seed weight also decreased by 6% in only the Swedish population but the difference was not significant (Fig S2). Overall, there is no differentiation between populations in total fitness in either treatment (Pop *p* = 0.781) but the Future treatment reduced fitness in the Swedish population marginally more than in the Italian population (Int *p* = 0.116) due to the decreased number of seeds per fruit. Despite being smaller plants overall, the Italian plants were able to match the fitness output of the Swedish plants by allocating almost twice as much biomass to reproduction in the Future treatment than Swedish plants (Fig S2; Pop *p* = 0.014).

**Figure 3.**
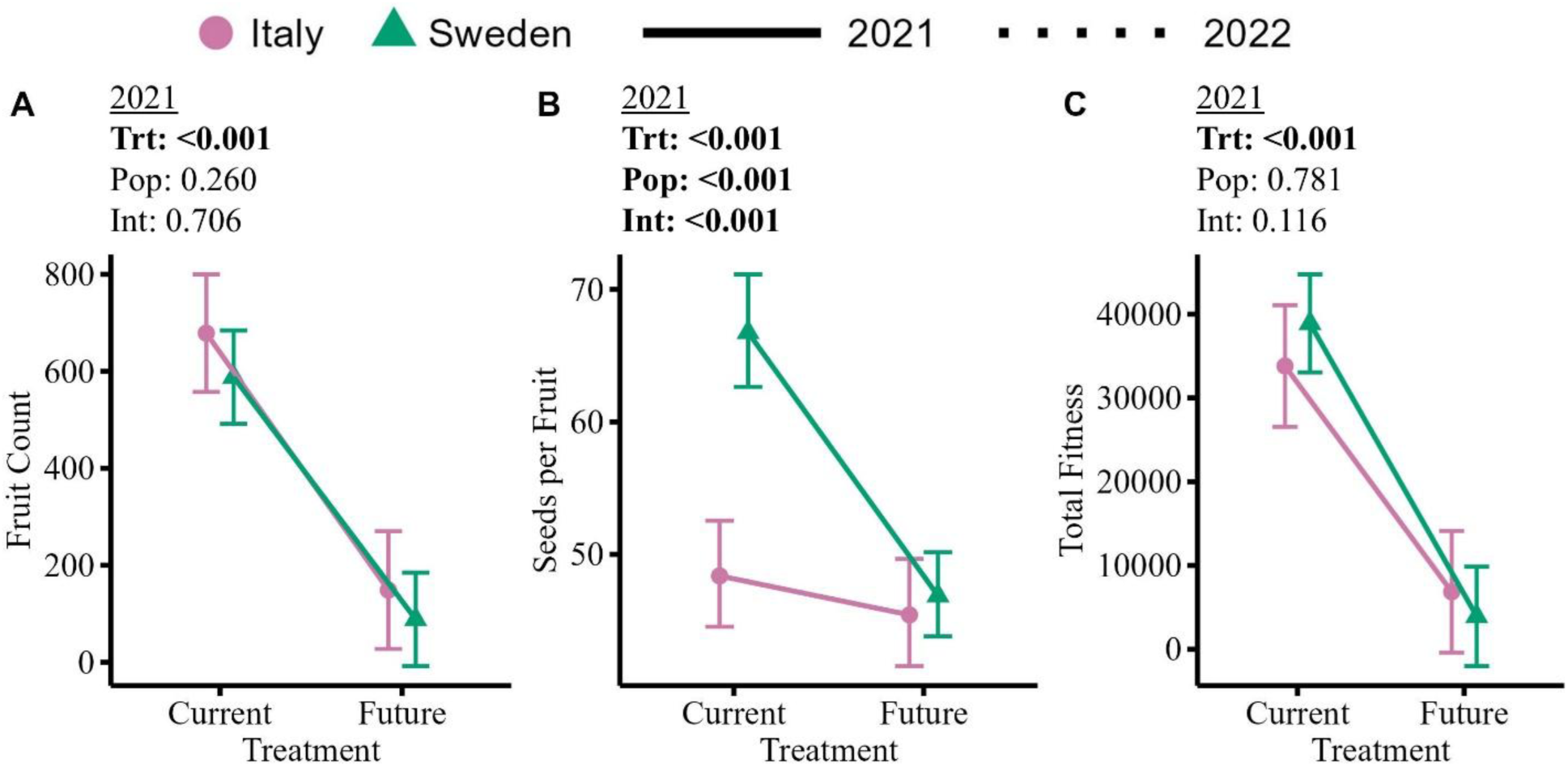
Heat and drought greatly decrease fitness for both populations. Each panel displays estimated marginal means for Sweden (triangles) and Italy (circles) with 95% confidence intervals. Lines connecting means illustrate plasticity; fitness was only measured in 2021 (solid lines). Statistics above each panel show *p*-values for Anova fixed effects. In fruit count (A) and total fitness (C), plants that produced no fruits are included but seeds per fruit (B) only includes plants that produced at least 1 fruit. **Alt Text:** Reaction norms labelled A through C comparing fruit count, seeds per fruit, and total fitness between populations and treatments from 2021. Going from the current treatment to the future treatment, both populations greatly decrease in fruit count and total fitness. Seeds per fruit decreases only for Sweden.

## DISCUSSION

### Adding heat to drought does not disrupt differentiation for drought strategy

Through replicated common garden experiments, we confirmed that *A. thaliana* from Sweden and Italy are genetically differentiated to avoid and escape drought, respectively, even if drought and heat are experienced concurrently. The Swedish population evolved in a climate with periodic drought during the plant life cycle; our results indicate it avoids drought through delayed flowering, greater biomass investment, and maintaining relative leaf water content (i.e., not wilting) when water availability is low (Figs 1-2). In contrast, the Italian population is adapted to an environment with a terminal drought; it escapes drought by flowering early and investing less in stress tolerant leaves (Figs 1-2). Our results use additional genotypes per population grown in heat and drought to confirm drought strategies identified through prior studies of *A. thaliana* with flowering time and water use efficiency in both the field and growth chambers (Ågren and Schemske 2012; Grillo *et al*. 2013; Dittmar *et al*. 2014; Mojica *et al*. 2016; Ågren *et al*. 2017; Oakley *et al*. 2023). Additionally, our results fit more broadly within a latitudinal gradient in resource-use strategy among European *A. thaliana* where populations from southern latitudes, like Italy, have a short lifespan with high SLA and northern populations, like Sweden, have a long lifespan and low SLA (Estarague et al. 2022).

Our results support prior work that the Swedish population avoids drought, but the absence of cold in both of our treatments complicates our conclusions. First, the same pathways involved in drought avoidance can also be involved in cold tolerance as both increase cellular protective compounds to avoid dehydration (Kong and Henry 2016; Suprasanna *et al*. 2016). Genotypes of *A. thaliana* from northern Europe had higher survival in both a cold treatment and a hot and dry treatment (Estarague *et al*. 2022). Our simulated winter stayed well above freezing, but cold tolerance is an important aspect of local adaptation of these populations (Oakley *et al*. 2018, 2023; Lee *et al*. 2024). We interpret lower SLA and greater shoot biomass as indicating greater vegetative investment by Sweden in drought avoidant leaves that are less prone to water loss. Those same trait means can also indicate cold tolerance so we cannot separate past selection in the native Swedish environment on drought avoidance from congruent selection on cold tolerance. Second, flowering in *A. thaliana* is controlled by a vernalization requirement in some populations (Michaels and Amasino 2000). While both populations are ecological winter annuals, the Italian population does not require vernalization to flower while the Swedish population does (Grillo *et al*. 2013). The difference in flowering time we observed between the populations is more similar to differences in flowering time observed in the field at the Italian site where temperatures are above freezing rather than the Swedish site where temperatures are below freezing for multiple weeks and there is little difference in days to flowering between the two populations (Ågren *et al*. 2017). Thus, our observed difference in flowering time may represent genetic differentiation for a vernalization requirement prior to flowering rather than represent genetic differentiation for drought strategy. However, the fact that the Italian genotypes flower earlier than the Swedish genotypes in both of our treatments is consistent with field experiments and a prior chamber study that did include temperatures below freezing to simulate a Swedish winter. Both studies identified that Italian alleles at flowering time quantitative trait loci cause earlier flowering (Dittmar *et al*. 2014; Ågren *et al*. 2017).

### Locally adapted populations respond to heat and drought stress with similar plasticity

We expected the Swedish population would be more plastic than the Italian population for most traits because Sweden has a more variable environment during the plant life cycle (Mojica *et al*. 2016) and plant populations from more variable precipitation regimes are more plastic for drought response traits (Gianoli and González-Teuber 2005). This was confirmed in prior work where the Swedish population was more plastic in response to drought (Mojica *et al*. 2016) and heat (Stewart *et al*. 2016). However, our results do not support this. Plasticity of both populations is in the direction expected of drought avoidance (i.e., generally in the direction of or more extreme than the Sweden mean phenotype in the Current treatment). Plasticity in the avoidant direction is not wholly unexpected because avoidance is favored when plants experience moderate water limitation during their life cycle and *A. thaliana* is generally considered avoidant (Verslues and Juenger 2011). Only two traits show genetic differentiation for plasticity: SLA and root-to-shoot ratio. Thus, drought strategy may contribute to the local adaptation of these populations, but both populations respond to the environmental cues of heat and drought with similar plasticity. This conserved plasticity is in agreement with prior work showing plasticity to mild drought stress is consistent across *A. thaliana* genotypes though there is variation in magnitude (Clauw *et al*. 2015).

Stomatal density was unexpectedly plastic in both populations. Our stomatal density values are two to five times greater than the stomatal density reported in other chamber studies using *A. thaliana* populations from the native range (Vile *et al*. 2012; Lorts and Lasky 2020) but about 33% less than transgenic genotypes developed to increase stomatal density (Tanaka *et al*. 2013). Vile *et al*. (2012) found stomatal density was not plastic between treatments similar to ours though it plastically decreased in heat and increased in drought compared to the cool, well-watered treatment. However, we observed plasticity to increase stomatal density in the Future treatment relative to the Current treatment and greater stomatal density in the Swedish population than in the Italian population. Sweden and Italy may have differentiation for stomatal density from evolving in climates with different drought regimes, but the contrast between our results and Vile *et al*. (2012) in stomatal density magnitude and plasticity even in the same species and similar chamber environments indicates that an underlying trait more directly involved in photosynthesis, such as stomatal conductance, may be more informative about plant responses to drought.

Further work should characterize if the observed plasticity is adaptive in both populations; our chamber study was limited in simulating field environments, and the sample sizes were too small to robustly test for adaptive plasticity (max sample size = 27 individuals but only 13 genotypes per population). Given the genetic differentiation for drought strategy between the populations, the conserved plasticity may be adaptive in one population but neutral or even maladaptive in the other. For example, decreased SLA in heat and drought is expected of drought avoidant plants. Decreasing SLA could be adaptive plasticity for the Swedish population but maladaptive for the Italian population. Understanding if plasticity in the Swedish population is adaptive will help predictions of population survival in future climates.

### Neither population had a fitness advantage in either treatment

The fitness of the Swedish population was not higher than that of the Italian population in our Current treatment, likely because our winter temperatures were well above freezing. In the field, there was no difference in total fitness measured by fruits per seedling planted between the Swedish and Italian population at the Swedish field site in only one of eight years (2010; Ågren and Schemske 2012; Ågren *et al*. 2013; Oakley *et al*. 2023), and if total fitness is measured by seeds per seedling planted, the Swedish population outperforms the Italian population in 2010 (Ellis *et al*. 2021). Cold tolerance is an important aspect of local adaptation between these populations (Oakley *et al*. 2018, 2023; Lee *et al*. 2024) and local adaptation of *A. thaliana* across the Eurasian range (Monroe *et al*. 2016). However, climate change is warming winter temperatures (IPCC 2021), so studies with warmer winter air temperatures like ours may help predict the impact of climate change on local adaptation and the efficacy of conservation efforts such as assisted gene flow (reviewed in (Anderson 2023)).

Natural gene flow is limited by dispersal and local adaptation to non-climate factors like soil properties and daylength (Corlett and Westcott 2013; Ni and Vellend 2024). Human-mediated migration of populations within or beyond their current ranges, particularly poleward or upward in elevation, may counteract an adaptation lag and prepare populations for future climates (Hewitt *et al*. 2011; Aitken and Whitlock 2013). Prior work with *A. thaliana* seeds collected between the 1930s and 2002 from populations across Europe identified an adaptation lag to warmer temperature almost 20 years ago; genotypes from locations historically warmer than their common garden sites had higher average fitness than even local genotypes suggesting the fitness optimum had already shifted (Wilczek *et al*. 2014). We thus expected Italy to have higher fitness in our Future treatment. However, the Swedish and Italian populations had the same mean fitness in our Future treatment, suggesting pure Italian genotypes are unlikely to outperform Swedish genotypes in a hotter and drier Swedish environment. However, primarily Swedish genotypes with introgression from Italy could outperform the native genotypes in a future climate. While a major concern about assisted gene flow, particularly in primarily self-fertilizing species, is a lack of introgression between resident and introduced populations (Wadgymar *et al*. 2015; Wadgymar and Weis 2017), there can be outcrossing in natural *A. thaliana* populations (Bomblies *et al*. 2010). The introduction of genetic variation for selection to act on through crosses between the Swedish and Italian populations may be crucial for the success of assisted gene flow in this system. The recombinant inbred lines developed with a parent from each population and grown at field sites already show transgressive variation for fitness (Ågren *et al*. 2013; Oakley *et al*. 2023) and could be used to further investigate hypotheses of assisted gene flow.

## CONCLUSIONS

We identified genetic differentiation between locally adapted populations of *A. thaliana* from Sweden and Italy that show Sweden avoids drought and Italy escapes drought. Despite this, both populations were plastic in the direction expected of drought avoidance when grown in a hotter and drier Swedish climate. If this evolutionarily conserved plasticity is adaptative remains unknown. Our hot and dry treatment greatly decreased fitness of both populations, raising concerns about population persistence under abiotic stress and the efficacy of assisted gene flow as a conservation strategy.

## SUPPLEMENTARY DATA

Referenced supplemental tables and figures will be made available online through the journal. All raw data and code will be made available through Zenodo upon manuscript acceptance.

## AUTHOR CONTRIBUTIONS

SFB, JKC, and EBJ contributed to idea generation. SFB managed the project. SFB conducted the 2021 experiment and VN conducted the 2022 experiment with mentorship from SFB and JKC. SFB and VN conducted initial analyses and wrote manuscript drafts. SFB finalized analyses and writing with input from JKC and EBJ.

## ACKNOWLEDGEMENTS

We thank Drs. Jon Ågren and Doug Schemske for sharing seeds. We thank Dr. Ågren as well as members of the Josephs and Conner labs for their constructive comments that greatly improved this manuscript. We thank the Michigan State Growth Chamber Facility for technical support.

