## Supplemental Information for "Differentiation for drought strategy but conserved plasticity to heat and drought in locally adapted populations of *Arabidopsis thaliana*"

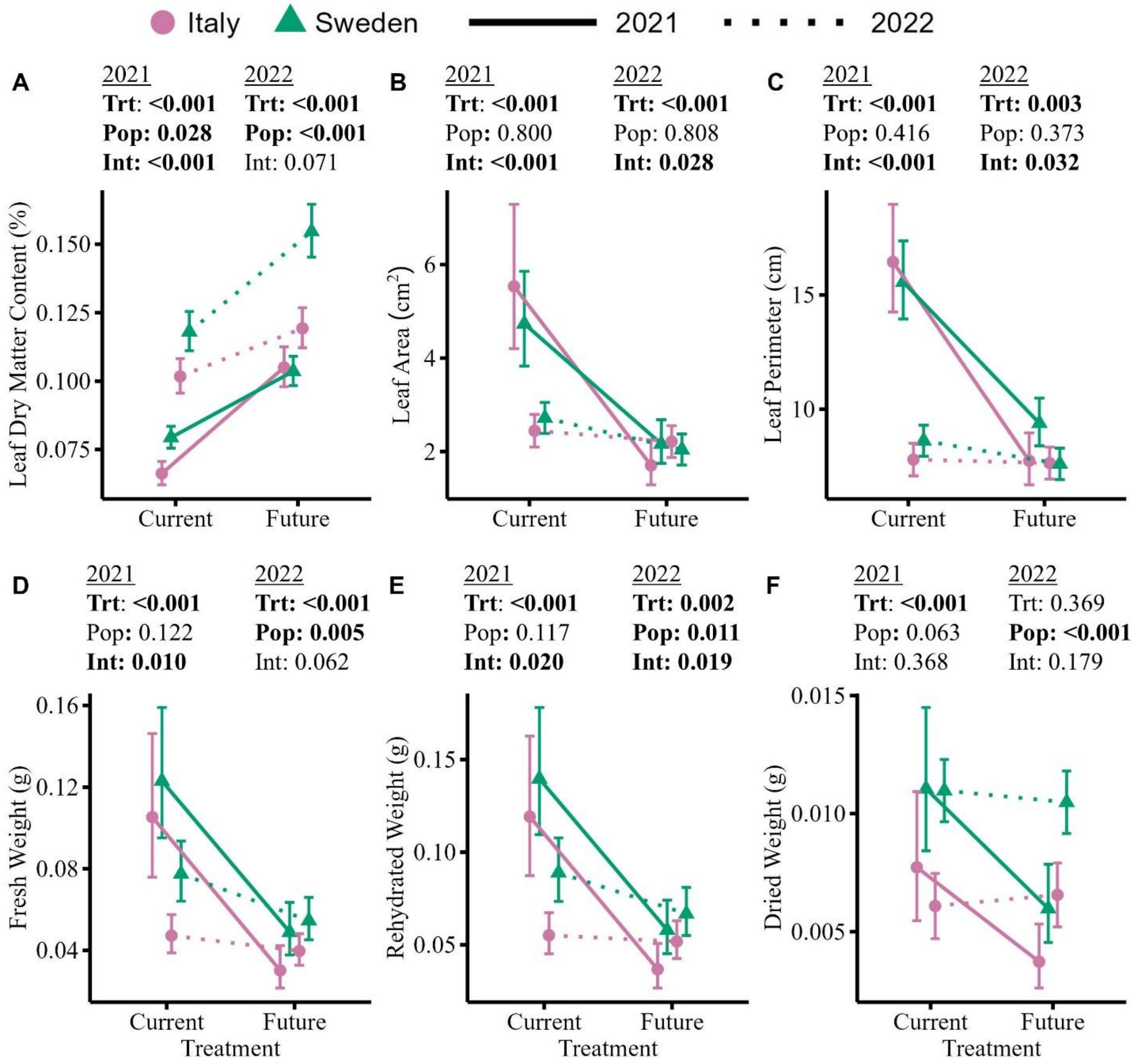


**Figure S1. Reaction norms for leaf dry matter content and components of specific leaf area and relative water content measured at bolting (2022) or flowering (2021).** Each panel displays estimated marginal means for Sweden (triangles) and Italy (circles) with 95% confidence intervals. Lines connecting means illustrate plasticity; the 2021 experiment is connected with solid lines and the 2022 experiment with dotted lines. Statistics above each panel show p-values for Anova fixed effects. Leaf dry matter content, fresh weight, and rehydrated weight were log transformed for statistical analysis in both experiments. Leaf area, leaf perimeter, and dry weight were log transformed in 2021 only.

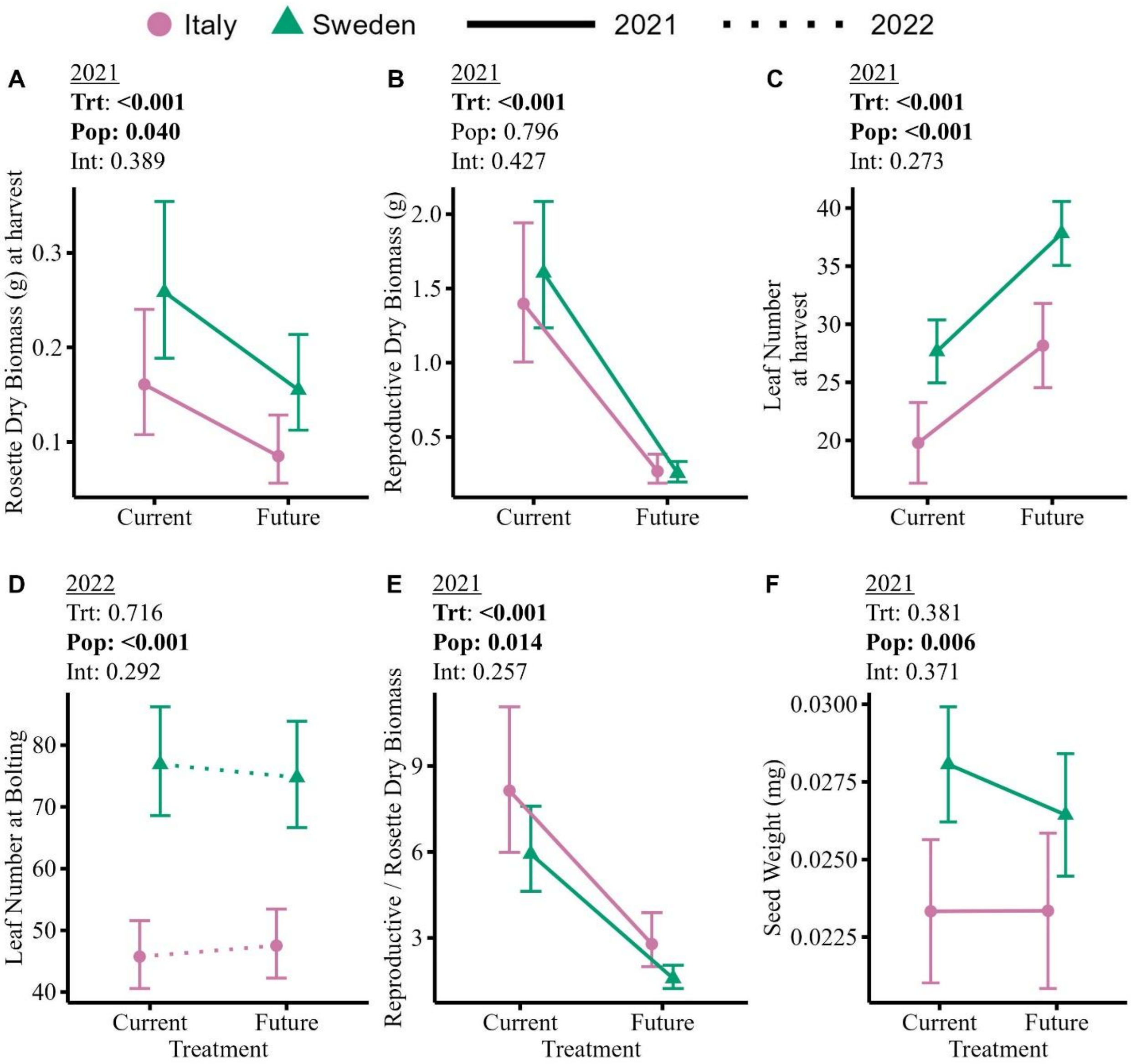


**Figure S2. Reaction norms for biomass, leaf number, and seed weight.** Each panel displays estimated marginal means for Sweden (triangles) and Italy (circles) with 95% confidence intervals. Lines connecting means illustrate plasticity; the 2021 experiment is connected with solid lines and the 2022 experiment with dotted lines. Statistics above each panel show p-values for Anova fixed effects. Rosette biomass, reproductive biomass and the ratio of reproductive:rosette biomass were log transformed for analysis. Note the difference in the y axis between panel C and D; rosettes in the 2022 experiment developed axillary meristem growth during the vegetative phase which greatly increased leaf number.

**Table S1.** Temperatures (°C) and soil moisture (%) for treatments in the 2021 experiment with their corresponding dates from the Sweden field site. Temperatures are written as Day Temperature:Night Temperature. Day lengths are denoted as hours:minutes.

| Week | Current | | Future | | Day Length | Simulated Dates |
|---|---|---|---|---|---|---|
| | Temp | Soil Moisture | Temp | Soil Moisture | | |
| 1 | 16:13 | 45 | 20:17 | 25 | 15:45 | Aug 1 - 23 |
| 2 | 14:11 | 45 | 18:15 | 25 | 14:15 | Aug 24 – Sept 15 |
| 3 | 8:6 | 45 | 12:10 | 25 | 12:00 | Sept 16 – Oct 8 |
| 4 | 8:6 | 45 | 10:9 | 25 | 9:30 | Oct 9 – 31 |
| 5 | 10:8 | 45 | 10:8 | 25 | 7:00 | Nov 1 – Dec 5 |
| 6 | 10:8 | 45 | 10:8 | 25 | 5:00 | Dec 6 – Jan 7 |
| 7 | 10:8 | 45 | 10:8 | 25 | 7:00 | Jan 8 – Feb 11 |
| 8 | 10:8 | 45 | 10:8 | 25 | 10:00 | Feb 12 – Mar 16 |
| 9 | 10:8 | 45 | 10:8 | 25 | 13:30 | Mar 17 – Apr 20 |
| 10 | 8:6 | 45 | 10:7 | 10 | 15:45 | Apr 21 – 27 |
| 11 | 8:6 | 45 | 12:7 | 10 | 16:30 | Apr 28 – May 4 |
| 12 | 9:6 | 45 | 13:10 | 10 | 17:00 | May 5 – 11 |
| 13 | 10:6 | 45 | 14:7 | 7.5 | 17:45 | May 12 – 18 |
| 14 | 9:6 | 45 | 13:10 | 7.5 | 18:30 | May 19 – 25 |
| 15 | 9:6 | 45 | 13:10 | 7.5 | 19:00 | May 26 – Jun 1 |
| 16 | 12:9 | 45 | 16:13 | 7.5 | 19:30 | June 2 – 8 |
| 17 | 10:9 | 45 | 14:13 | 7.5 | 19:45 | June 9 – 15 |
| 18 | 15:11 | 45 | 19:15 | 7.5 | 20:00 | June 16 – 22 |
| 19 | 14:11 | 45 | 18:15 | 7.5 | 20:00 | June 23 – 29 |
| 20 | 18:12.5 | 45 | 22:16.5 | 7.5 | 19:45 | June 30 – July 6 |
| 21 | 20:13.5 | 45 | 24:17.5 | 7.5 | 19:15 | July 7 – 13 |
| 22 | 16:13 | 45 | 20:17 | 7.5 | 18:45 | July 14 – 20 |
| 23 | 16:13 | 45 | 20:17 | 7.5 | 18:45 | July 14 – 20 |
| 24 | 16:12 | 45 | 20:16 | 7.5 | 18:15 | July 21 – 27 |
| 25 | 18:13 | 45 | 22:17 | 7.5 | 17:30 | July 28 – Aug 3 |

**Table S2.** Temperatures (°C) and soil moisture (%) for treatments in the 2021 experiment with their corresponding dates from the Sweden field site if applicable. Temperatures are written as Day Temperature:Night Temperature. Day lengths are denoted as hours:minutes.

| Week | Current | | Future | | Day Length | Simulated Dates |
|---|---|---|---|---|---|---|
| | Temp | Soil Moisture | Temp | Soil Moisture | | |
| 1 | 16:13 | 45 | 16:13 | 7.5 | 16:00 | Fall |
| 2 | 16:13 | 45 | 16:13 | 7.5 | 16:00 | Fall |
| 3 | 16:13 | 45 | 16:13 | 7.5 | 16:00 | Fall |
| 4 | 10:8 | 45 | 10:8 | 7.5 | 7:00 | Nov 1 – Dec 5 |
| 5 | 10:8 | 45 | 10:8 | 7.5 | 5:00 | Dec 6 – Jan 7 |
| 6 | 10:8 | 45 | 10:8 | 7.5 | 7:00 | Jan 8 – Feb 11 |
| 7 | 10:8 | 45 | 10:8 | 7.5 | 10:00 | Feb 12 – Mar 16 |
| 8 | 10:8 | 45 | 10:8 | 7.5 | 13:30 | Mar 17 – Apr 20 |
| 9 | 10:6 | 45 | 14:10 | 7.5 | 17:00 | May 5 – 11 |
| 10 | 10:6 | 45 | 14:10 | 7.5 | 17:45 | May 12 – 18 |
| 11 | 10:6 | 45 | 14:10 | 7.5 | 18:30 | May 19 – 25 |
| 12 | 10:6 | 45 | 14:10 | 7.5 | 19:00 | May 26 – Jun 1 |
| 13 | 12:9 | 45 | 16:13 | 7.5 | 19:30 | June 2 – 8 |
| 14 | 10:9 | 45 | 14:13 | 7.5 | 19:45 | June 9 – 15 |
| 15 | 15:11 | 45 | 19:15 | 10 | 20:00 | June 16 – 22 |
| 16 | 14:11 | 45 | 18:15 | 10 | 20:00 | June 23 – 29 |
| 17 | 18:12.5 | 45 | 22:16.5 | 10 | 19:45 | June 30 – July 6 |
| 18 | 20:13.5 | 45 | 24:17.5 | 10 | 19:15 | July 7 – 13 |

**Table S3.** Anova results for all traits in supplemental. Columns denote the trait used as the response variable in the model, the predictor model terms, and then for the 2021 and 2022 experiments the sum of squares (always matches the mean square), the denominator degrees of freedom, *F* value, and *p* value. Numerator degrees of freedom is always 1 for treatment, population, and their interaction; it is always 2 for block. Symbols illustrate *p* value: † < 0.1; * <0.05; ** < 0.01; *** < 0.001.

| Experiment | | 2021 | | | | 2022 | | | |
|---|---|---|---|---|---|---|---|---|---|
| Trait | Term | SumSq | DenDF | *F* | *p* | SumSq | DenDF | *F* | *p* |
| Log Fresh Weight (g) | Treatment | 4.18578 | 58.7753 | 309.4517 | 4.29E-25*** | 0.239980546 | 107.187 | 18.48769 | 3.78E-05*** |
| | Population | 0.036572 | 14.50807 | 2.703716 | 0.121602133 | 0.147691964 | 12.37973 | 11.37793 | 0.005317677** |
| | Interaction | 0.095456 | 58.7753 | 7.057006 | 0.010148993* | 0.046084358 | 106.7838 | 3.55026 | 0.062255752† |
| | Block | Not applicable | | | | 0.005245664 | 106.8209 | 0.202058 | 0.8173587 |
| Log Rehydrated Weight (g) | Treatment | 3.755336 | 58.45017 | 275.6137 | 8.56E-24*** | 0.127457966 | 106.1692 | 9.792431 | 0.002262794** |
| | Population | 0.038005 | 14.07761 | 2.789297 | 0.116971279 | 0.11655334 | 12.39128 | 8.954643 | 0.01087761** |
| | Interaction | 0.077542 | 58.45017 | 5.69101 | 0.020312618* | 0.073363894 | 105.8146 | 5.636453 | 0.019393437* |
| | Block | Not applicable | | | | 0.007523902 | 105.8474 | 0.289026 | 0.74958205 |
| Log Dried Weight (g) | Treatment | 1.604446 | 57.94126 | 114.4465 | 2.43E-15*** | 3.27E-06 | 108.1626 | 0.815269 | 0.368573414 |
| | Population | 0.057241 | 14.0165 | 4.083066 | 0.062847498† | 0.000112541 | 13.12876 | 28.03886 | 0.00014035*** |
| | Interaction | 0.011528 | 57.94126 | 0.822284 | 0.368269003 | 7.34E-06 | 107.8033 | 1.827712 | 0.179227673 |
| | Block | Not applicable | | | | 7.21E-06 | 107.8425 | 0.898709 | 0.410121325 |
| Log Leaf Area | Treatment | 3.282806 | 57.10278 | 315.5779 | 6.26E-25*** | 4.958156459 | 107.0874 | 15.81636 | 0.000127119*** |
| | Population | 0.000691 | 13.64002 | 0.066425 | 0.80046236 | 0.019382622 | 11.87248 | 0.06183 | 0.807875864 |
| | Interaction | 0.133158 | 57.10278 | 12.80059 | 0.000714675*** | 1.56420631 | 106.7394 | 4.989767 | 0.027582949* |
| | Block | Not applicable | | | | 0.349518675 | 106.7904 | 0.557477 | 0.574309716 |

| Experiment | | 2021 | | | | 2022 | | | |
|---|---|---|---|---|---|---|---|---|---|
| Trait | Term | SumSq | DenDF | *F* | *p* | SumSq | DenDF | *F* | *p* |
| Log Leaf Dry Matter Content | Treatment | 0.484606 | 66.43312 | 275.252 | 2.57E-25*** | 0.251919037 | 120 | 45.9357 | 4.85E-10*** |
| | Population | 0.010224 | 16.8207 | 5.807098 | 0.027708223* | 0.247549383 | 120 | 45.13892 | 6.52E-10*** |
| | Interaction | 0.034755 | 66.43312 | 19.74027 | 3.44E-05*** | 0.018187604 | 120 | 3.316384 | 0.071083851† |
| | Block | Not applicable | | | | 0.036858467 | 120 | 3.360444 | 0.038017401* |
| Log Dried Rosette Mass (g) | Treatment | 1.148173 | 57.45863 | 62.87137 | 8.56E-11*** | Not measured | | | |
| | Population | 0.092911 | 14.88376 | 5.087609 | 0.039589781* | | | | |
| | Interaction | 0.013737 | 57.45863 | 0.752188 | 0.389391959 | | | | |
| Log Dry Repro Mass (g) | Treatment | 10.93581 | 67.38359 | 224.1077 | 4.08E-23*** | Not measured | | | |
| | Population | 0.003355 | 19.77693 | 0.068764 | 0.795856743 | | | | |
| | Interaction | 0.031159 | 67.38359 | 0.638547 | 0.427045133 | | | | |
| Leaf Number at bolting | Treatment | Not measured | | | | 0.000771902 | 109.1628 | 0.132778 | 0.716273328 |
| | Population | | | | | 0.263839288 | 14.20356 | 45.38424 | 8.84E-06*** |
| | Interaction | | | | | 0.006529829 | 108.8304 | 1.123227 | 0.291571195 |
| | Block | | | | | 0.007348982 | 108.8674 | 0.632067 | 0.53343129 |
| Leaf Number at harvest | Treatment | 1628.54 | 62.36899 | 132.9939 | 4.16E-17*** | Not measured | | | |
| | Population | 230.8288 | 16.5638 | 18.85052 | 0.00046862*** | | | | |
| | Interaction | 14.98058 | 62.36899 | 1.223382 | 0.272946244 | | | | |
| Log Repro:Rosette Biomass | Treatment | 5.069443 | 67.96255 | 123.2345 | 6.48E-17*** | Not measured | | | |
| | Population | 0.289278 | 23.48758 | 7.032148 | 0.014106575* | | | | |
| | Interaction | 0.053673 | 67.96255 | 1.304761 | 0.257353762 | | | | |
| | Block | Not applicable | | | | | | | |

| Experiment | | 2021 | | | | 2022 | | | |
|---|---|---|---|---|---|---|---|---|---|
| Trait | Term | SumSq | DenDF | *F* | *p* | SumSq | DenDF | *F* | *p* |
| Average Seed Weight | Treatment | 1.20E-05 | 59.26482 | 0.779703 | 0.38079977 | Not measured | | | |
| | Population | 0.00017 | 12.5741 | 11.01171 | 0.005780445** | | | | |
| | Interaction | 1.25E-05 | 59.26482 | 0.812051 | 0.371162029 | | | | |
